# MetaDome: Pathogenicity analysis of genetic variants through aggregation of homologous human protein domains

**DOI:** 10.1101/509935

**Authors:** Laurens Wiel, Coos Baakman, Daan Gilissen, Joris A. Veltman, Gerrit Vriend, Christian Gilissen

## Abstract

The growing availability of human genetic variation has given rise to novel methods of measuring genetic tolerance that better interpret variants of unknown significance. We recently developed a novel concept based on protein domain homology in the human genome to improve variant interpretation. For this purpose we mapped population variation from the Exome Aggregation Consortium (ExAC) and pathogenic mutations from the Human Gene Mutation Database (HGMD) onto Pfam protein domains. The aggregation of these variation data across homologous domains into meta-domains allowed us to generate base-pair resolution of genetic intolerance profiles for human protein domains.

Here we developed MetaDome, a fast and easy-to-use web service that visualizes meta-domain information and gene-wide profiles of genetic tolerance. We updated the underlying data of MetaDome to contain information from 56,319 human transcripts, 71,419 protein domains, 12,164,292 genetic variants from gnomAD, and 34,076 pathogenic mutations from ClinVar. MetaDome allows researchers to easily investigate their variants of interest for the presence or absence of variation at corresponding positions within homologous domains. We illustrate the added value of MetaDome by an example that highlights how it may help in the interpretation of variants of unknown significance. The MetaDome web server is freely accessible at https://stuart.radboudumc.nl/metadome.

## Introduction

The continuous accumulation of human genomic data has spurred the development of new methods to interpret genetic variants. There are many freely available web servers and services that facilitate the use of these data by non-bioinformaticians. For example, the ESP Exome Variant Server (Fu et al., 2012; NHLBI GO Exome Sequencing Project (ESP), 2011) and the Genome Aggregation Database (gnomAD) browser (Karczewski et al., 2017; Lek et al., 2016) help locate variants that occur frequently in the general population. These services are used for the interpretation of unknown variants based on the assumption that variants occurring frequently in the general population are unlikely to be relevant for patients with Mendelian disorders (Amr et al., 2016). There are also methods that derive information from these large human genetic databases. For example genetic intolerance, which is commonly used to interpret variants of unknown significance by assessing whether variants stand out because they occur in regions that are genetically invariable in the general population (Ge et al., 2016; Gussow et al., 2016). Examples of such methods are RVIS (Petrovski, Wang, Heinzen, Allen, & Goldstein, 2013) and subRVIS (Gussow, Petrovski, Wang, Allen, & Goldstein, 2016). The strongest evidence for the pathogenicity of a genomic variant comes from the presence of that variant in any of the clinically relevant genetic variant databases such as the Human Gene Mutation Database (HGMD) (Stenson et al., 2017) or the public archive of clinically relevant variants (ClinVar) (Landrum et al., 2016). These databases are gradually growing in the amount of validated pathogenic information.

Another way to provide evidence for the pathogenicity of a genomic variant is to observe the effect of that variant in homologous proteins across different species. Mutations at corresponding locations in homologous proteins are found to result in similar effects on protein stability (Ashenberg, Gong, & Bloom, 2013). Finding homologous proteins is one the key applications of BLAST (Altschul, Gish, Miller, Myers, & Lipman, 1990). Transferring information between homologous proteins is one of the oldest concepts in bioinformatics, and can be achieved by performing a multiple sequence alignment (MSA) and locating equivalent positions between the protein sequences. We have previously used this concept and showed that it also holds for homologous Pfam protein domain relationships within the human genome. We found that ~71-72% of all disease-causing missense variants from HGMD and ClinVar occur in regions translating to a Pfam protein domain and observed that pathogenic missense variants at equivalent domain positions are often paired with the absence of population-based variation and *vice versa* (Wiel, Venselaar, Veltman, Vriend, & Gilissen, 2017). By aggregating variant information over homologous protein domains, the resolution of genetic tolerance per position is increased to the number of aligned positions. Similarly, the annotation of pathogenic variants found at equivalent domain positions also assists the interpretation of variants of unknown significance. We realized that this type of information could be of great benefit to the genetics community and therefore developed MetaDome.

MetaDome is a freely available web server that uses our concept of ‘meta-domains’ (i.e. aggregated homologous domains) to optimally use the information from population-based and pathogenic variation datasets without the need of a bioinformatics intermediate. MetaDome is easy to use and utilizes the latest population datasets by incorporating the gnomAD and ClinVar datasets.

## Results

### Accessibility

The MetaDome web server is freely accessible at https://stuart.radboudumc.nl/metadome. MetaDome features a user-friendly web interface and features a fully interactive tour to get familiar with all parts of the analysis and visualizations.

All source code and detailed configuration instructions are available in our GitHub repository: https://github.com/cmbi/metadome.

### The underlying database: a mapping between genes and proteins

The MetaDome web server queries genomic datasets in order to annotate positions in a protein or a protein domain. Therefore, the server needs access to genomic positional information as well as protein sequence and protein domain information. The database maps GENCODE gene translations to entries in the UniProtKB/Swiss-Prot databank in a per-position manner and corresponding protein domains or genomic variation. With respect to our criteria to map gene translations to proteins (**Methods; creating the mapping database**), 42,116 of the 56,319 full-length protein-coding GENCODE Basic transcripts for 19,728 human genes are linked to 33,492 of the 42,130 Swiss-Prot human canonical or isoform sequences. Of the total 591,556 canonical and isoform sequences present in Swiss-Prot, 42,130 result from the Human species. The resulting mappings contain 32,595,355 unique genomic positions that are linked to 19,226,961 residues in Swiss-Prot protein sequences.

71,419 Pfam domains are linked to 30,406 of the Swiss-Prot sequences in our database. Of these Pfam domain instances, 5,948 are from a unique Pfam domain family and 3,334 of these families have two or more homologues and are therefore suitable for meta-domain construction. Thus, by incorporating every protein-coding transcript, instead of only the longest ones, we increase the previously 2,750 (Wiel et al., 2017) meta-domains to 3,334. These meta-domains, on average, consist of 16 human protein domain homologues with a protein sequence length of 158 residues. Table 1 summarizes the counting statistics for sequences, domains, etc.

**Table 1.** Statistics on the number of entries present in GENCODE, Swiss-Prot, and our mapping database.

| Database | What | # of entries |
| --- | --- | --- |
| GENCODE | Protein-coding genes | 20,345 |
| MetaDome | Protein-coding genes | 19,728 |
| GENCODE | Protein-coding transcripts | 57,005 |
| MetaDome | Protein-coding transcripts | 56,319 |
| Swiss-Prot | Canonical and isoform protein sequences | 591,556 |
| Swiss-Prot | Human canonical and isoform protein sequences | 42,130 |
| MetaDome | Gene translations identically mapped to a canonical or isoform protein sequence | 42,116 |
| MetaDome | Canonical and isoform protein sequences | 33,492 |
| MetaDome | Pfam protein domain regions | 71,419 |
| MetaDome | Unique Pfam protein domain families | 5,948 |
| MetaDome | Unique Pfam protein domain families with two or more within-human occurrences | 3,334 |
| MetaDome | Chromosome to protein position mappings | 70,261,143 |
| MetaDome | Unique chromosome positions | 32,595,355 |
| MetaDome | Unique residues (as part of a protein) | 19,226,961 |
| MetaDome | Unique protein sequences with at least one Pfam domain annotated | 30,406 |

### How to use the MetaDome web server

At the welcome page users are offered the option to start an interactive tour or start with the analysis. The navigation bar at the top is available throughout all web pages in MetaDome and allow for further navigation to the ‘About’, ‘Method’, ‘Contact’ page (Supp. Figure S1). The user can fill in a gene symbol in the ‘gene of interest’ field and is aided by an auto-completion to help you find your gene of interest more easily (Supp. Figure S2). Clicking the ‘Get transcripts’ fills all GENCODE transcripts for that gene in the dropdown box. Only the transcripts that are mapped to a Swiss-Prot protein can be used in the analysis, the others are displayed in grey (Supp. Figure S3).

Clicking the ‘Start Analysis’ button starts an extensive query to the back-end of the web server for the selected transcript. Firstly, all the mappings are retrieved for the transcript of interest. Secondly, the entire transcript is annotated with ClinVar and gnomAD single nucleotide variants (SNVs) and Pfam domains. Thirdly, if there are any Pfam domains suitable for meta-domain relations then all mappings for those regions are gathered and annotated with ClinVar and gnomAD variation (**methods; Composing a meta-domain**).

The web-page provided to the user as a result of the ‘Analyse Protein’ can best be explained using an example. Therefore, we have generated this result for gene *CDK13* for transcript ‘ENST00000181839.4’ (Figure 1). The result page features four main components that we will describe from top to bottom. Located at the top is the graph control field. Directly below the graph control is the landscape view of the protein. Below the landscape view, a schematic and interactive representation of the protein and an additional representation of the protein which controls the zooming option. Lastly, at the bottom of the page there is the list of selected positions. All of these components are interactive and the various functionalities are described in Table 2.

**Figure 1.**
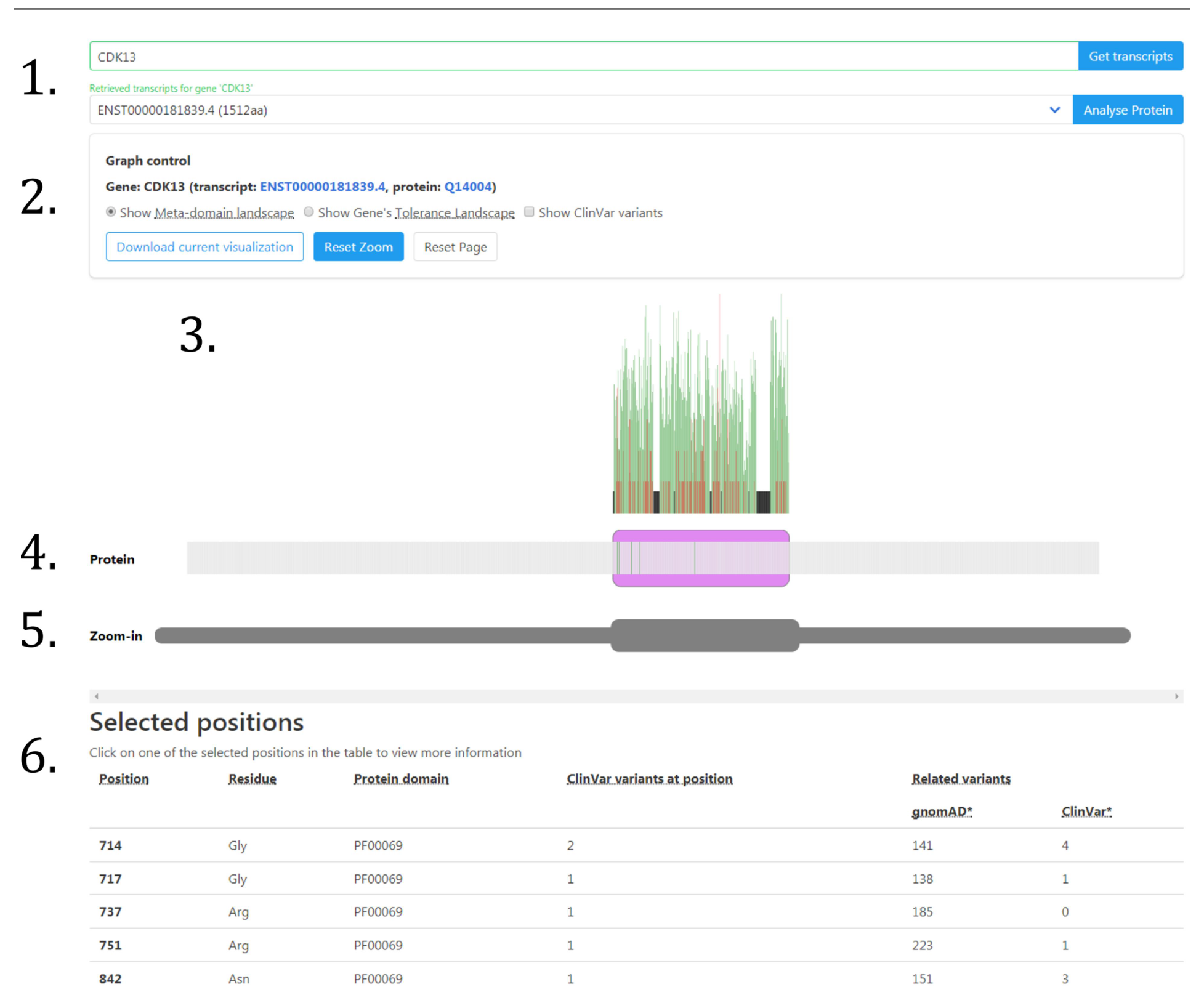
MetaDome web server result for the gene *CDK13*. The result provided by the MetaDome web server for the analysis of gene *CDK13* with transcript ENST00000181839.4, as provided in 1.). In 2.), there is additional information that the translation of this transcript corresponds to Swiss-Prot protein Q14004. Here also various alternative visualizations can be selected. The visualization starts by default in the ‘meta-domain landscape’, a mode selectable in the graph control in 2.). The landscapes are visualized in 3.), and in the meta-domain landscape the domain regions are annotated with missense variation counts found in homologous domains as bar plots. The schematic protein representation, located at 4.), is per-position selectable, and the domains are presented as purple blocks. Selected positions are highlighted in green. The ‘Zoom-in’ section at 5.) features a selectable greyed-out copy of schematic protein representation that can zoom-in on any part of the protein. Any selected positions are in the list of selected positions in 6.). Here more information can be obtained by clicking on one of these positions. A detailed description of the functionality of each component is described in Table 2.

**Table 2.**
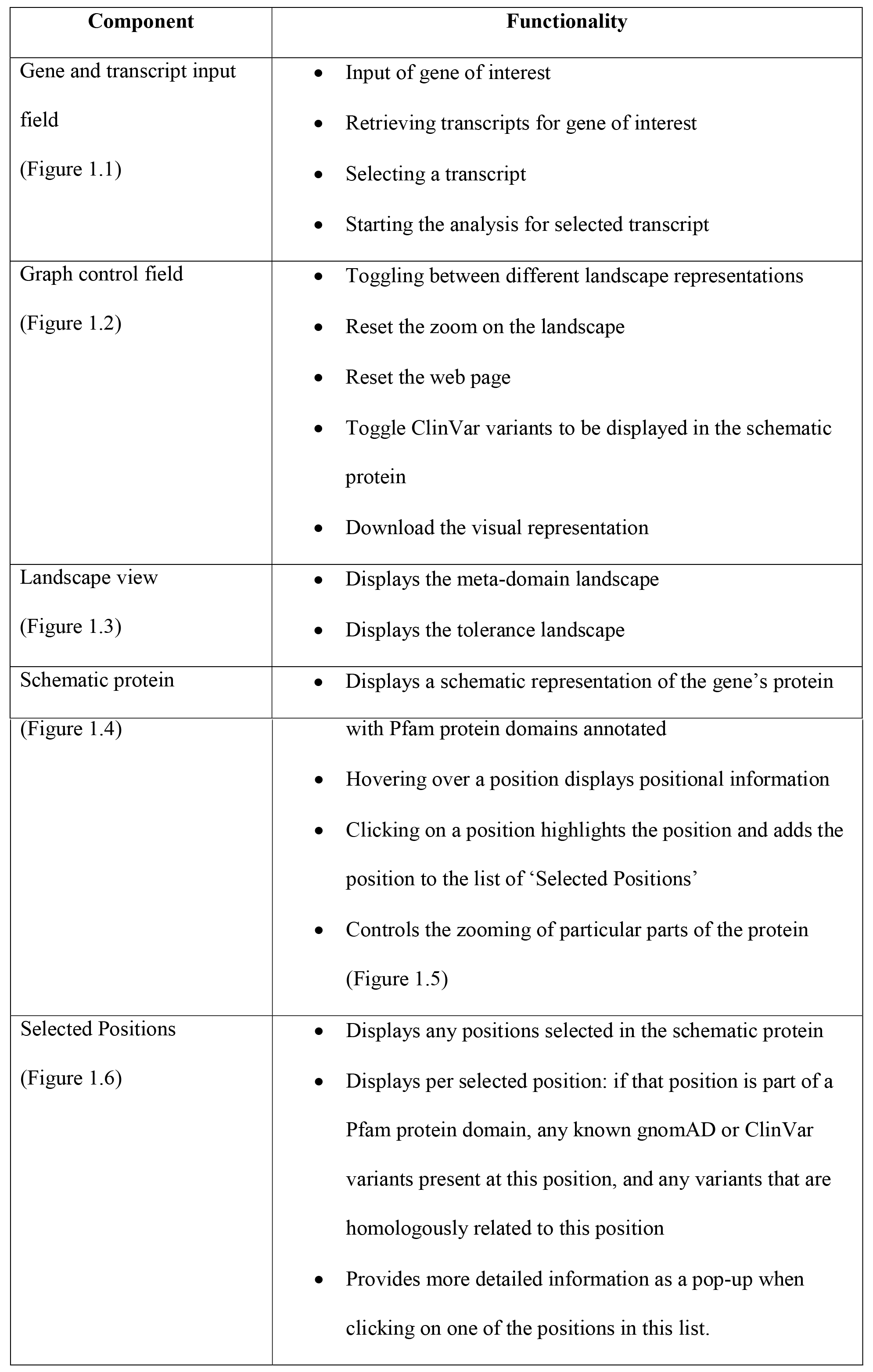
Descriptions of the various functionalities on the MetaDome result page.

Another way to use population-based variation in the context of the entire protein is via the tolerance landscape representation in MetaDome that can be selected in the graph control component (Figure 1.2). The tolerance landscape depicts a missense over synonymous ratio (also known as *K_a_/K_S_* or *d_N_/d_S_*) over a sliding window of 21 residues over the entirety of the protein of interest (e.g. calculated for ten residues left and right of each residue) based on the gnomAD dataset (**methods; Computing genetic tolerance and generating a tolerance landscape;** Figure 2A). Previously, the *d_N_/d_S_* metric has been used by others and us to measure genetic tolerance and predict disease genes (Ge, Kwok, & Shieh, 2015; Gilissen et al., 2014; Lelieveld et al., 2017), and it is suitable for measuring tolerance in regions within genes (Ge et al., 2016).

**Figure 2.**
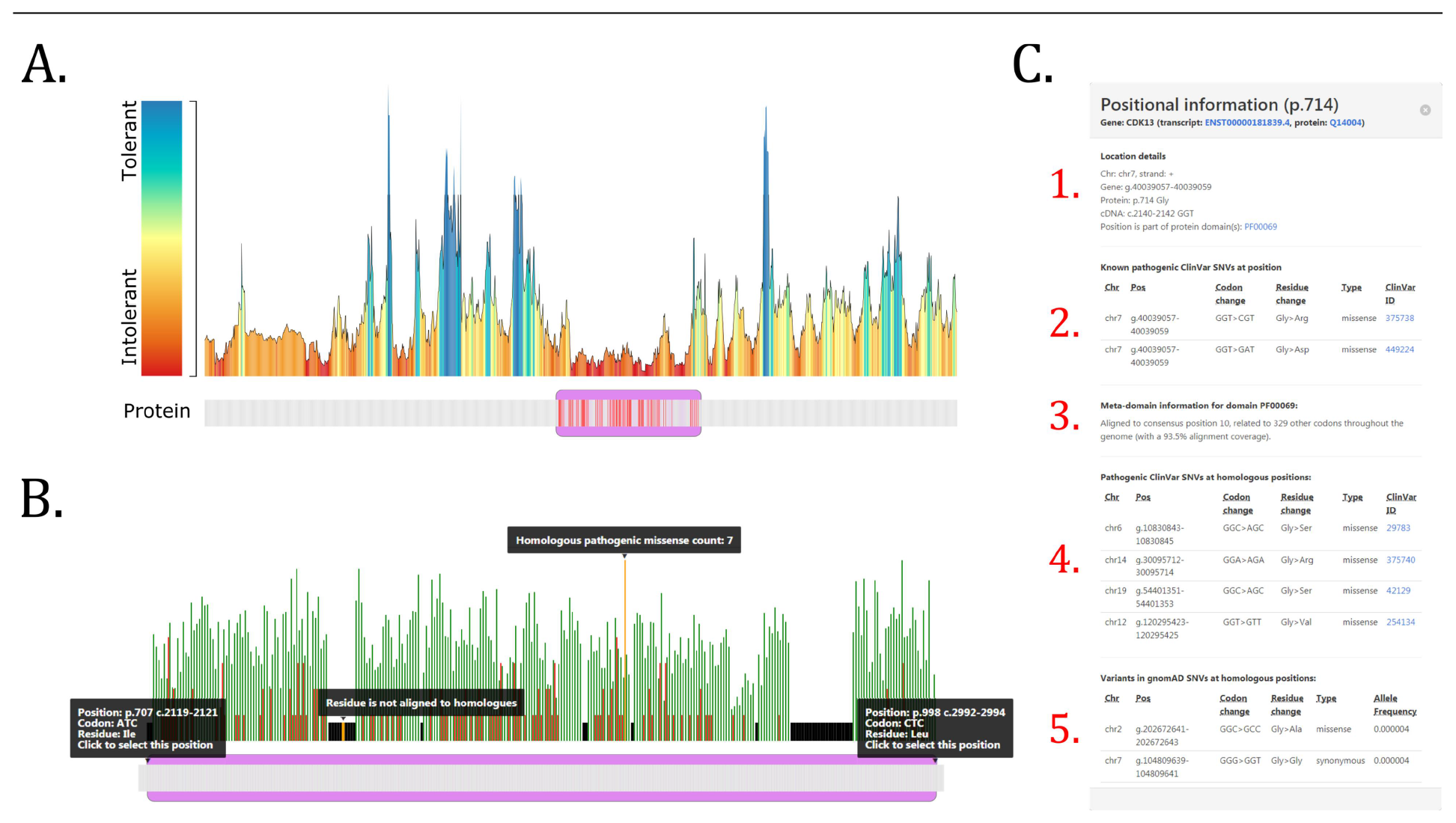
Examples of a MetaDome analysis for the gene *CDK13*. **A.)** The tolerance landscape depicts a missense over synonymous ratio calculated as a sliding window over the entirety of the protein (**methods; Computing genetic tolerance and generating a tolerance landscape**). The missense and synonymous variation are annotated from the gnomAD dataset and the landscape provides some indication of regions that are intolerant to missense variation. In this *CDK13* tolerance landscape the Pkinase Pfam protein domain (PF00069) in purple can be clearly seen as intolerant if compared to other parts in this protein. The red bars in the schematic protein representation correspond to pathogenic ClinVar variants found in this gene and in homologous protein domains. All of these variants are contained in the intolerant region of the landscape. **B.)** A zoom-in on the meta-domain landscape for *CDK13.* The Pkinase Pfam protein domain (PF00069) is located between protein positions 707 and 998 and annotated as a purple box in the schematic protein representation. The meta-domain landscape displays a deep annotation of the protein domain: the green (gnomAD) and red (ClinVar) bars correspond to the amount of missense variants found at aligned homologous positions. Unaligned positions are annotated as black bars. All of this information is displayed upon hovering over these various elements. **C.)** The positional information provides a detailed overview of a position from the ‘Selected Positions’ list, especially if that position is aligned to domain homologues. Here, for position p.Gly714 we can observe in 1.) the positional details for this specific protein position. In 2.) is any known pathogenic information for this position. We can observe here that for this position there are two known pathogenic missense variants. In 3.) meta-domain information is displayed and we can observe that p.Gly714 is aligned to consensus position 10 in the Pkinase Pfam protein domain and related to 329 other codons. This consensus position has an alignment coverage of 93.5% for the meta-domain MSA. There are also four pathogenic variants found in ClinVar on corresponding homologous positions as can be seen in 4.) and in 5.) there is an overview of all corresponding variants found in gnomAD.

### An example of using the MetaDome web server for variant interpretation

The MetaDome analysis result for *CDK13* (Figure 1) is the longest protein coding transcript for *CDK13* with a protein sequence length of 1,512 amino acids. In the resulting schematic protein representation we can observe the Pkinase Pfam protein domain (PF00069) between positions 707 and 998 as the only protein domain in this gene (Figure 2B). The Pkinase domain is highly prevalent throughout the human genome with as many as 779 homologous occurrences in human proteins, of which 353 are unique genomic regions. It is the 8th most occurring domain in our mapping database. The meta-domain landscape is the default view mode and shows any missense variation found in homologous domain occurrences throughout the human genome. Population-based (gnomAD) missense variation is displayed in green and pathogenic (ClinVar) missense variation is annotated in red bars, with the height of the bars depicting the number of variants found at each position (Figure 2B).

If the ‘Show ClinVar variants’ is toggled in the graph control field, we may observe that all known pathogenic information is highlight positions in the schematic protein in a red colour and for this example is located in the protein domain (Figure 2A). All ClinVar variants annotated to the gene are displayed this way in red. Additionally, any ClinVar variants that are present at homologous positions are also displayed in red. In total six known disease causing SNVs are present in the *CDK13* gene itself according to ClinVar, and these all fall within the Pkinase protein domain. All of these are missense variants. If we add variants found in homologous domains there are 64 positions with one or more reported pathogenic variants (**Supp Data S1**). Four of these positions overlap with the positions on which ClinVar variants were found in the gene itself and on position p.883 (Supp. Figure S4) we can observe a peak of eight missense variants annotated from other protein domains.

MetaDome helps to look in more detail to a position of interest. If we do this for protein position 714 (Figure 2C) in *CDK13* we find that it corresponds to consensus position 10 in the Pkinase domain (PF00069). At this position in *CDK13* there are two variants reported in ClinVar: p.Gly714Arg (ClinVar ID: 375738) submitted by (Sifrim et al., 2016), and p.Gly714Asp (ClinVar ID: 449224) submitted by GeneDX. The first is reported as a *de novo* variant and is associated to Congenital Heart Defects, Dysmorphic Facial Features, and Intellectual Developmental Disorder. For the second there is no associated phenotype provided. As MetaDome annotates variants reported at homologous positions, we can find even more information for this particular position. At the homologues aligned to this position we find a variant of identical change in *PRKD1:* p.Gly600Arg (ClinVar ID: 375740) reported as pathogenic and *de novo* in the same study (Sifrim et al., 2016). It is also associated to Congenital Heart Defects as well as associated to Ectodermal Dysplasia. There are three more reported pathogenic variants aligned to this position: MAK:p.Gly13Ser (ClinVar ID: 29783) associated to Retinitis Pigmentosa 62 (Özgül et al., 2011), PRKCG:p.Gly360Ser (ClinVar ID: 42129) associated to Spinocerebellar Ataxia Type14 (Klebe et al., 2005), and CIT:p.Gly106Val (ClinVar ID: 254134) associated to Microcephaly 17, primary, autosomal recessive (Özgül et al., 2011). These homologously related pathogenic variants and the severity of the associated phenotypes contributes to the evidence that this particular residue may be important at this position. Further evidence can be found from the fact that in human homologue domains this residue is extremely conserved. There are 330 unique genomic regions encoding for a codon aligned to this position (**Supp Data S2**). Only in the gene *PIK3R4* (ENST00000356763.3) does this codon encode for another residue than Glycine, namely a Threonine at position p.Thr35.

In the same way that we explored pathogenic ClinVar variation we can also explore the variation reported in gnomAD. In *CDK13* at protein position 714 there is no reported variant in gnomAD, but there are homologously related variations. There are 65 missense variants with average allele frequency of 1.24E-05 and 76 synonymous with average allele frequency 8.71E-03 and there is no reported nonsense variation (**Supp Data S1**).

When we inspect the tolerance landscape for *CDK13* (Figure 2A) we can see that all of the ClinVar variants (either annotated in *CDK13* or related via homologues) fall within the Pkinase Pfam protein domain (PF00069). In addition, the protein domain can clearly be seen as more intolerant to missense variation as compared to other parts of this protein, thereby supporting the ClinVar variants likely pathogenic role.

## Conclusion

The MetaDome web server combines resources and information from different fields of expertise (e.g. genomics and proteomics) in order to increase the power in analysing population and pathogenic variation by transposing this variation to homologous protein domains. Such a transfer of information is achieved by a per-position mapping between the GENCODE and Swiss-Prot databases. 79.4% of the Human Swiss-Prot protein sequences are of identical match to one or more of 42,116 GENCODE transcripts. This means that 25.7% of the GENCODE transcriptions differ in mRNA but translate to the same Swiss-Prot protein sequence. GENCODE previously reported that this is due to alternative splicing, of which a substantial proportion only affect untranslated regions (UTRs) and thus have no impact on the protein-coding part of the gene (Harrow et al., 2006).

MetaDome is especially informative if a variant of interest falls within a protein domain that has homologues. This is highly likely as 43.6% of the positions in the MetaDome mapping database are part of a homologous protein domain. Pathogenic missense variation is also highly likely to fall within a protein domain as we previously observed for 71% of HGMD and 72% of ClinVar pathogenic missense variants (Wiel et al., 2017). By aggregating variation over protein domain homologues via MetaDome, the resolution of genetic tolerance at a single base-pair is increased. Furthermore, we can obtain variation that could disrupt the functionality of a protein domain, as annotated throughout the entire human genome, which may potentially be disease-causing. It should be noted, that by aggregating genetic variation in this way the specific context such as haplotype information or interactions with other proteins may be lost. Aggregation via meta-domains only encapsulates general biological or molecular functions attributed to the domain. Nonetheless, we believe MetaDome can be used to better interpret variants of unknown significance through the use of meta-domains and tolerance landscapes as we have shown in our example.

As more genetic data accumulates in the years to come, MetaDome will become more and more accurate in predictions of intolerance at the base-pair level and the meta-domain landscapes will become even more populated with variation found in homologue protein domains. We can imagine many other ways of integrating this type of information to be helpful for variant interpretation. Future directions for the MetaDome web server could lead to machine learning empowered variant effect prediction, or visualization of the meta-domain information in a protein 3D structure.

## Methods

### Software architecture of MetaDome

MetaDome is developed in Python v3.5.1 (Rossum & Drake, 2010) and makes use of the Flask framework v0.12.4 (Ronacher, 2010) for the web service part which communicates between the front-end, the back-end, and the database. The software architecture (Supp. Fig S5) follows the Domain-driven design paradigm (Evans, 2004). The code is open source and can be found at our GitHub repository: https://github.com/cmbi/metadome. Detailed instructions on how to deploy the MetaDome web server can be found there too.

To ensure MetaDome can be deployed to any environment and provide a high degree of modularity, we have containerized the application via Docker v17.12.1 (Hykes, 2013). We use docker-compose v1.17.1 to ensure that different containerized aspects of the MetaDome server can work together. The following aspects are containerized to this purpose: 1.) The Flask application, 2.) a PostgreSQL v10 database wherein the mapping database is stored, 3.) a Celery v4.2.0 task queue management system to facilitate the larger tasks of the MetaDome web-based user requests, 4.) a Redis v4.0.11 for task result storage, and 5.) RabbitMQ v3.7 to mediate as a task broker between client and workers. For a full overview of the docker-compose architecture we refer to Supp. Fig. S6.

The visualization medium of the MetaDome web server is a fully interactive and responsive HTML web page. This page is generated by the Flask framework and the navigation aesthetics are made using the CSS framework Bulma v0.7.1 (Thomas, Potiekhin, Lauhakari, Shah, & Berning, 2018). The visualizations of the various landscapes and the schematic protein are created with JavaScript, JQuery v3.3.1, and the D3 Framework v4.13.0 (Bostock, Ogievetsky, & Heer, 2011).

### Datasets of population and disease-causing genetic variation

MetaDome makes use of single nucleotide variants (SNVs) from population and clinically relevant genetic variation databases. Population variation was obtained from the gnomAD r2.0.2 VCF file by selecting all synonymous, nonsense, and missense variants that meet the PASS filter criteria. Variants meeting the PASS criteria are considered to be true variants (Lek et al., 2016). The variants in the VCF file from ClinVar release 2018 05 03 with disease-causing (Pathogenic) status are used as the disease-causing SNVs in MetaDome.

### Creating the mapping database

MetaDome stores a complete mapping between genomic, protein positions, and all domain annotations (Supp. Fig. S7) in a PostgreSQL relational database (PostgreSQL Global Development Group, 1996). This mapping is auto-generated and stored in the PostgreSQL database by the MetaDome web server upon the first run. The genomic positions consist of each chromosomal position in the protein-coding transcripts of the GENCODE release 19 GRCh37.p13 Basic set (Harrow et al., 2012). The protein positions correspond to protein sequence positions in the UniProtKB/Swiss-Prot Release 2016_09 databank entries for the human species (Boutet et al., 2016). These mappings are created with Protein-Protein BLAST v2.2.31+ (Camacho et al., 2009) for each protein-coding translation in the GENCODE Basic set to human canonical and isoform Swiss-Prot protein sequences. We exclude sequences that do not start with a start codon (i.e. ATG encoding for methionine), or end with a stop codon. We checked if the cDNA sequence of the transcripts match the GENCODE translation via Biopython’s translate function (Cock et al., 2009), if they are not identical than these are excluded too. The global information on the transcript (e.g. identifiers, sequence length) is registered in the database in the table ‘genes’ and, for each Swiss-Prot entry with an identical sequence match, the global information is stored in the table ‘proteins’.

Next, for each identical match between translation and Swiss-Prot sequence a ClustalW2 v2.1 (Larkin et al., 2007) alignment is made between these two sequences. Each nucleotide’s genomic position is mapped to the protein position and stored in the ‘mappings’ table. Each entry in mapping represents a single nucleotide of a codon and is linked to the corresponding entry in the ‘genes’ and ‘proteins’ table (i.e. the corresponding GENCODE translation, transcription and Swiss-Prot sequence).

Each Swiss-Prot sequence in the database is annotated via InterProScan v5.20-59.0 (Finn et al., 2017) for Pfam-A v30.0 protein domains (Finn et al., 2016) and the results are stored in the ‘interpro_domains’ table. After the construction of the database is finished, all meta-domain alignments can be constructed.

### Composing a meta-domain

Meta-domains consist of homologous Pfam protein domain instances that are annotated using InterproScan. Meta-domains consist of at least two homologous domains. MSAs are made using a three step process. 1.) Retrieve all sequences for the domain instances, 2.) Retrieve the Pfam HMM corresponding to the Pfam identifier annotated by InterproScan, and 3.) Use HMMER 3.1b2 (Finn et al., 2015) to align the sequences from the first step. The resulting Stockholm format MSA files can be inspected with alignment visualization software like Jalview (Waterhouse, Procter, Martin, Clamp, & Barton, 2009). In this Stockholm formatted file, all columns that correspond to the domain consensus represent the same homologous positions.

These Stockholm files are retrieved by the MetaDome web server when a user request meta-domain information for a position of their interest. Upon retrieval of this Stockholm file, the mapping database is used to obtain the corresponding genomic positions for each residue. These genomic positions are subsequently used to retrieve corresponding gnomAD or ClinVar variation.

### Computing genetic tolerance and generating a tolerance landscape

The non-synonymous over synonymous ratio, or *d_N_/d_g_* score, is used to quantify genetic tolerance. This score is based on the observed (obs) missense and synonymous variation in gnomAD (*missense_obs_ and synonymous_obs_*). This score is corrected for the sequence composition by taking into account the background (bg) of possible missense and synonymous variants based on the codon table (*missense_bg_ and synonymous_bg_*):

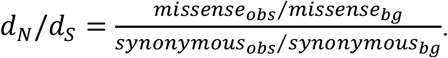

The tolerance landscape computes this ratio as a sliding window of size 21 (i.e. ten residues before and ten after the residue of interest) over the entirety of the gene’s protein. The edges (e.g. start and end) are therefore a bit noisy as they are not the result of averaging over a full length window.

## Acknowledgements

This work was in part financially supported by grants from the Netherlands Organization for Scientific Research (916-14-043 to C.G. and 918-15-667 to J.A.V.), and from the Radboud Institute for Molecular Life Sciences, Radboud university medical center (R0002793 to G.V.). We thank Hanka Venselaar for her critically reading of the manuscript.

## URLs

MetaDome web service: https://stuart.radboudumc.nl/metadome

GENCODE: https://www.gencodegenes.org/

InterPro: https://www.ebi.ac.uk/interpro/

Pfam: https://pfam.xfam.org/

gnomAD: http://gnomad.broadinstitute.org/

ClinVar: https://www.ncbi.nlm.nih.gov/clinvar/

GitHub repository: https://github.com/cmbi/metadome

## Supplemental Figures

**Supp Fig. S1.**
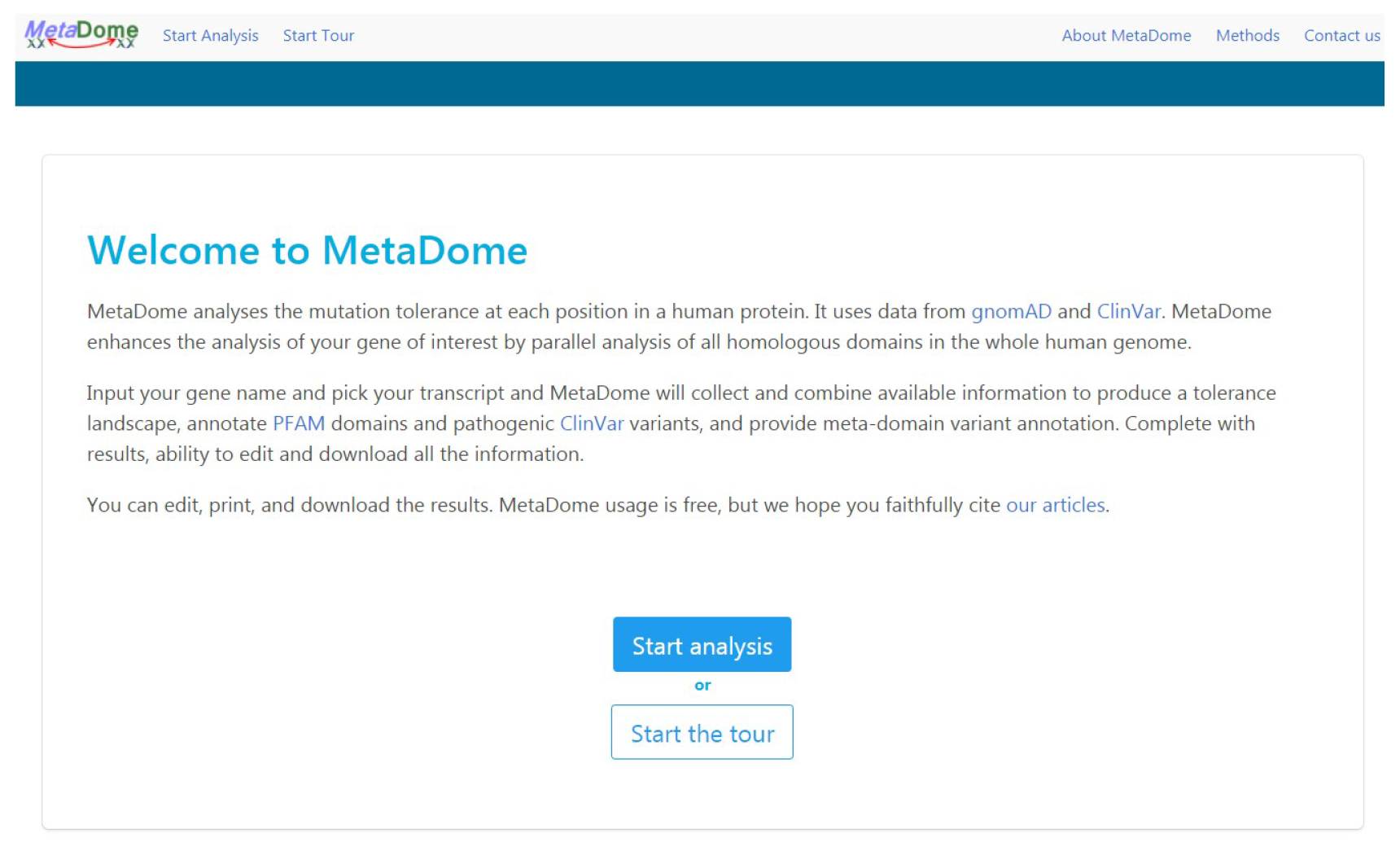
Welcome page. The welcome page of MetaDome is the entry point to the rest of the web server. Here the navigation bar is located at the top that eases navigation throughout the rest of the web pages. From here the user can start the interactive tour or an analysis.

**Supp Fig. S2.**
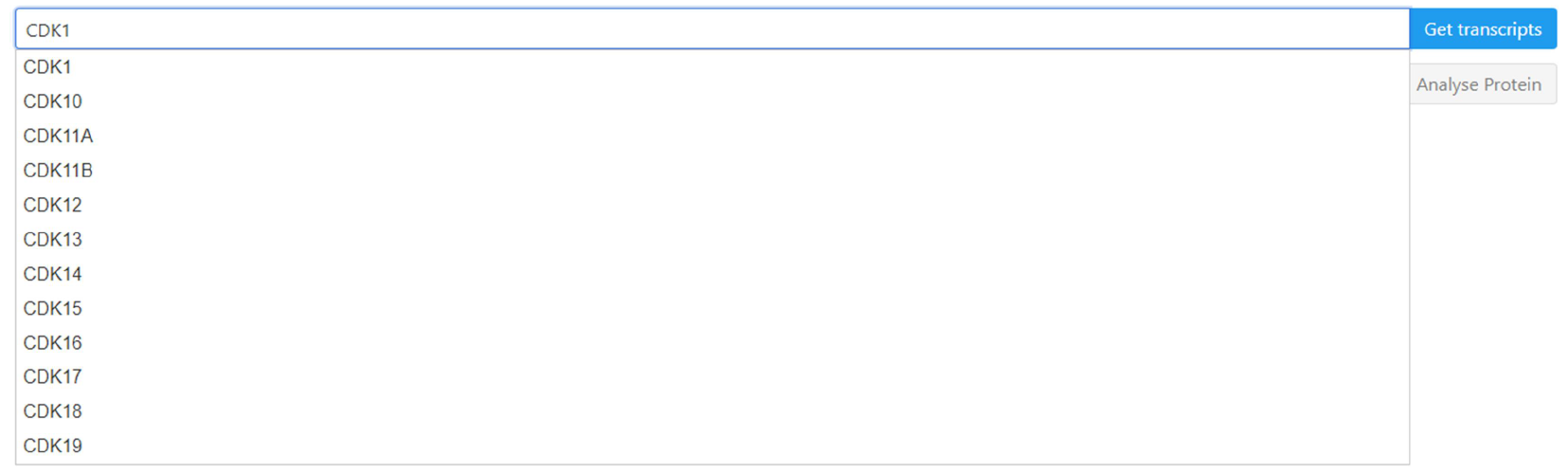
Gene symbol auto-completion. The auto-completion of gene symbols is based on all gene symbols present in the mapping database. The auto-completion will start once the user has entered three characters in the input field and is interactive (e.g. you can click on the gene that is of your interest.

**Supp Fig. S3.**
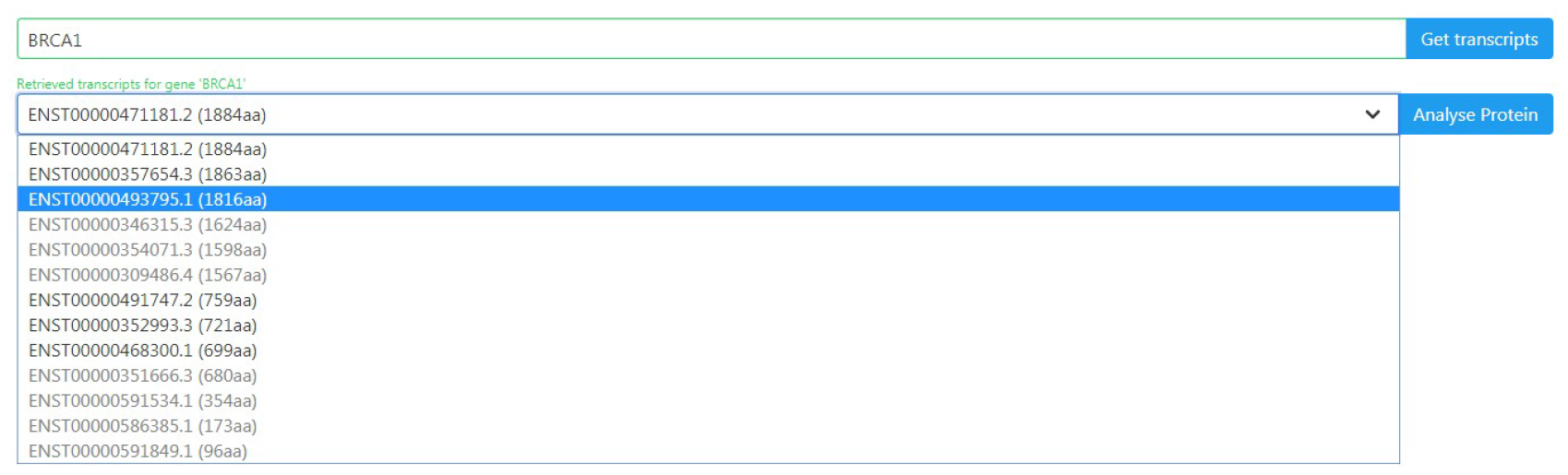
Dropdown box containing transcript identifiers. The result of clicking the ‘Get transcripts’ button for the *BRCA1* gene. Here a success message is displayed in green, which means that the gene is present in the database. The resulting transcripts are ordered by amino acid (aa) length and the longest transcripts are at the top. The greyed-out transcripts are not suitable for further analysis in the web server.

**Supp Fig. S4.**
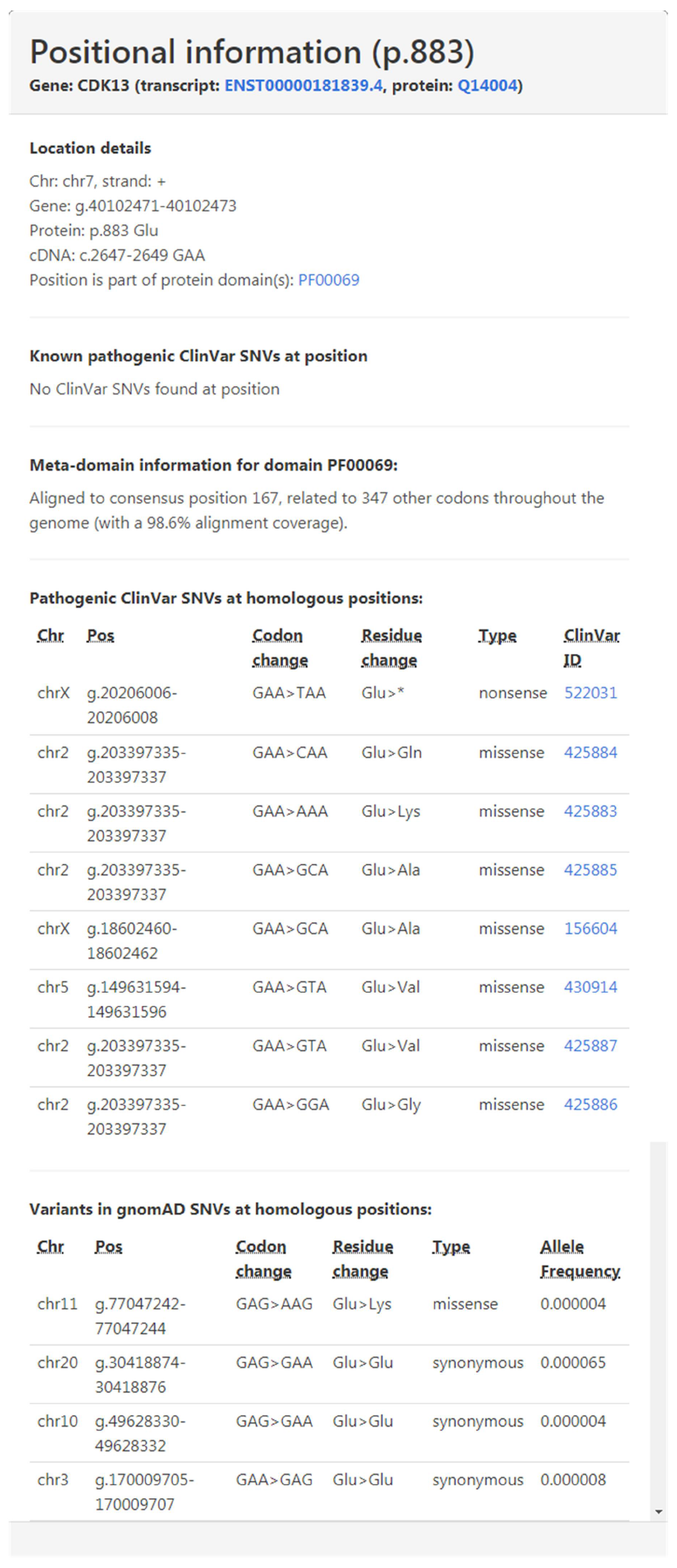
Dropdown box containing transcript identifiers. The positional information overview provides detailed positional information for position p.Glu883. We can observe that it is aligned to consensus position 167 in the Pkinase Pfam protein domain. This consensus position has 98.6% alignment coverage. On this position no ClinVar pathogenic variants are found in *CDK13*, but on homologous positions there are eight pathogenic ClinVar variants annotated.

**Supp Fig. S5.**
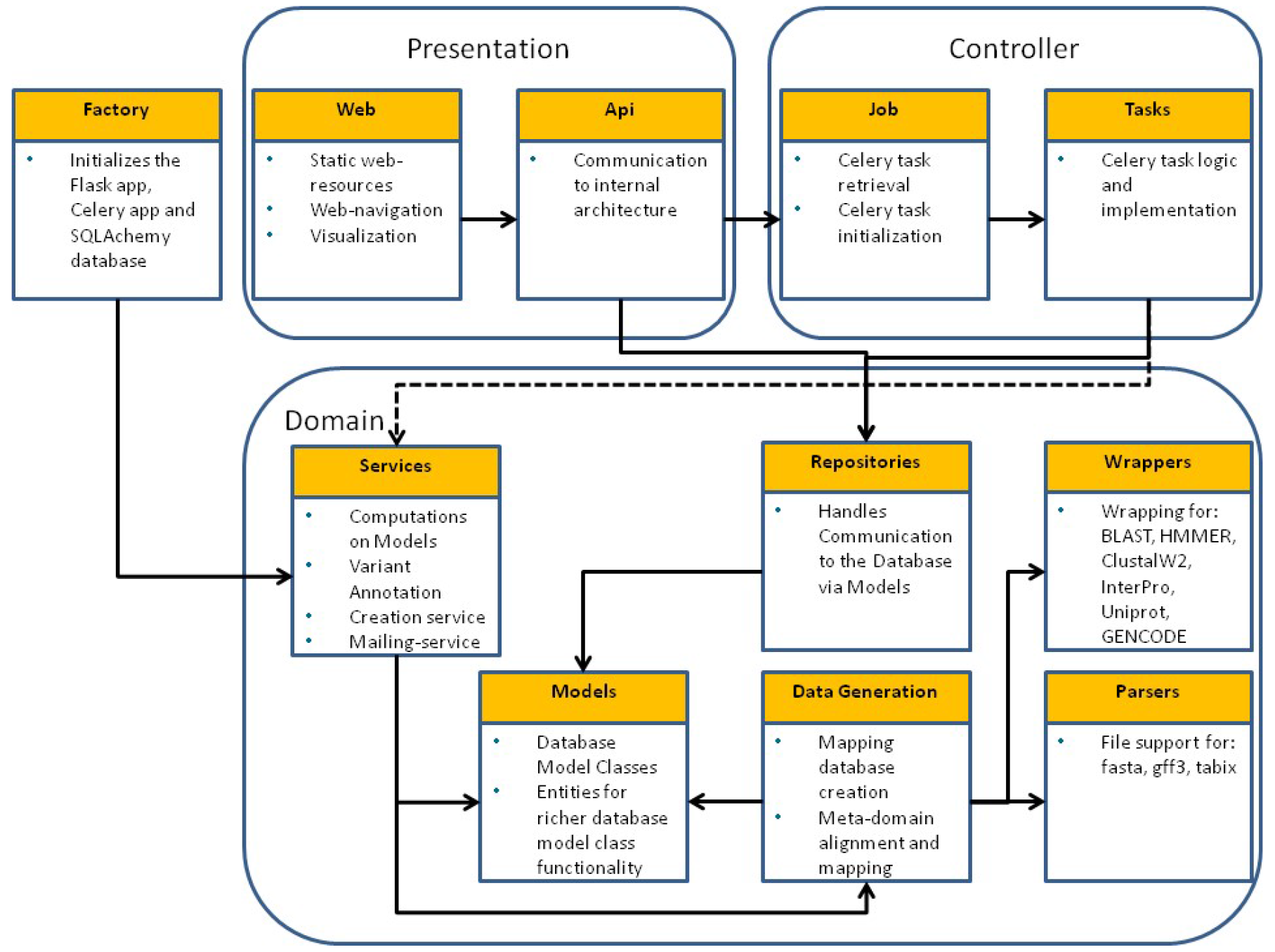
The software architecture of the MetaDome web server. A high-level overview of the software architecture of the MetaDome web server. The architecture follows closely the Domain-driven design paradigm. In this architecture we make use of three distinct layers with separated responsibilities. The Presentation layer is responsible for the user interface and external communication handling. This layer takes care of all the visualizations in the web interface and any user input that needs further validation or that requires starting a celery job. In the controller layer all communication is handled from the MetaDome server to the Celery task queue. Job here instantiates and checks the status of jobs in the Celery queue that are initialized from Tasks. The Domain layer contains all the domain-knowledge of the MetaDome application. Here Repositories is the outside connection for approaching the internal database, so no domain knowledge is needed by the Presentation layer. The services take care of various computations on the data from Models and contains functionality to create the database via Data Generation. The Factory is outside the layers, but initializes the configuration to run the MetaDome server (e.g. connections to the PostgreSQL and Celery Docker containers and start of the Flask web service).

**Supp Fig. S6.**
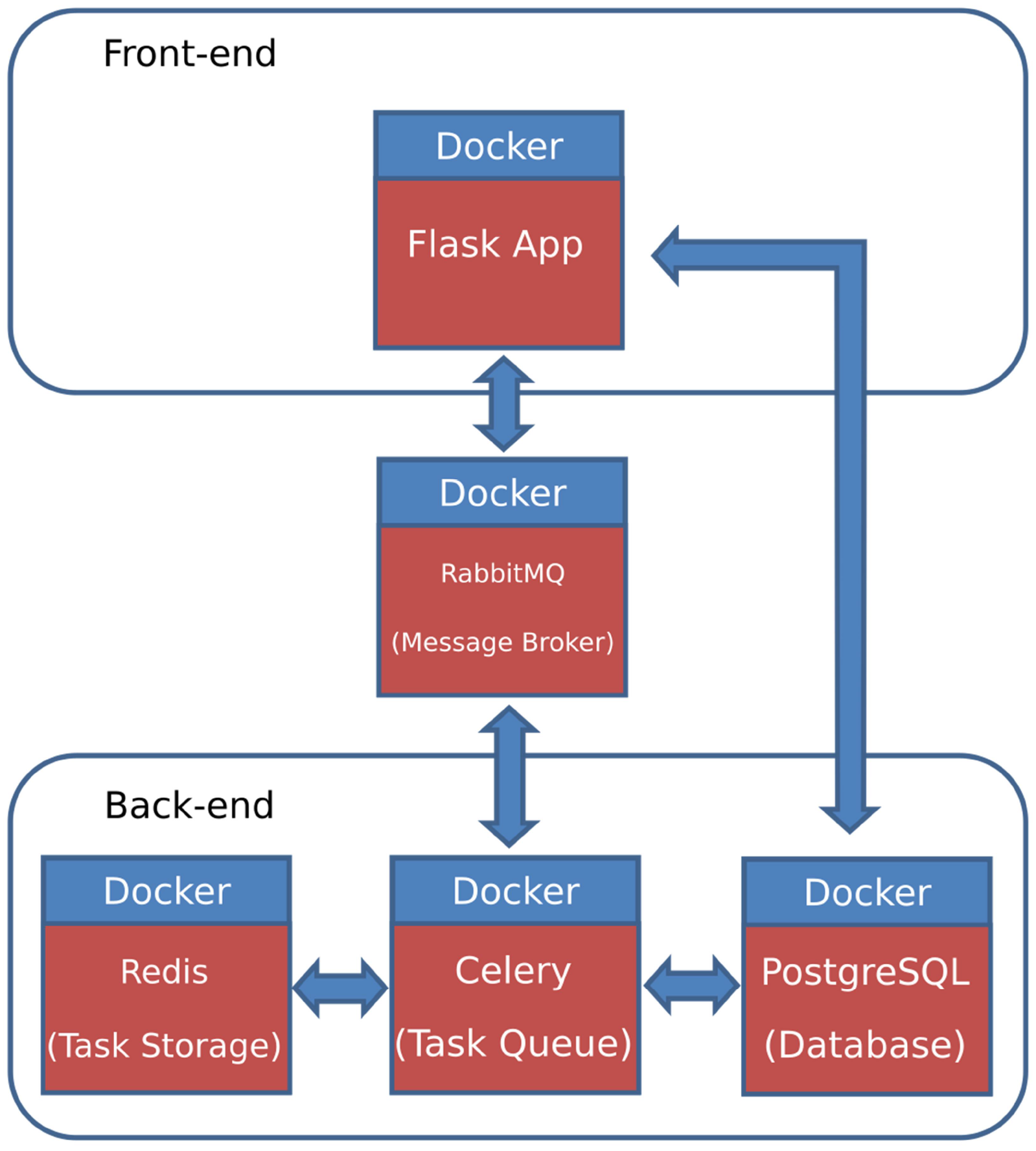
The internal docker network architecture of the MetaDome web server. Metadome makes use of several docker containers that work jointly via docker-compose to provide various degrees of functionality. The Flask App docker container is responsible for hosting the MetaDome visual interface and at the first run it will create the mapping database. The Flask docker container communicates via the RabbitMQ service, which serves as a message broker, to the Celery docker container. This Celery container keeps track of the large tasks and the results of those tasks are temporarily stored in the Redis server. The PostgreSQL container houses the database containing the mapping between GENCODE transcripts and Swiss-Prot.

**Supp Fig. S7.**
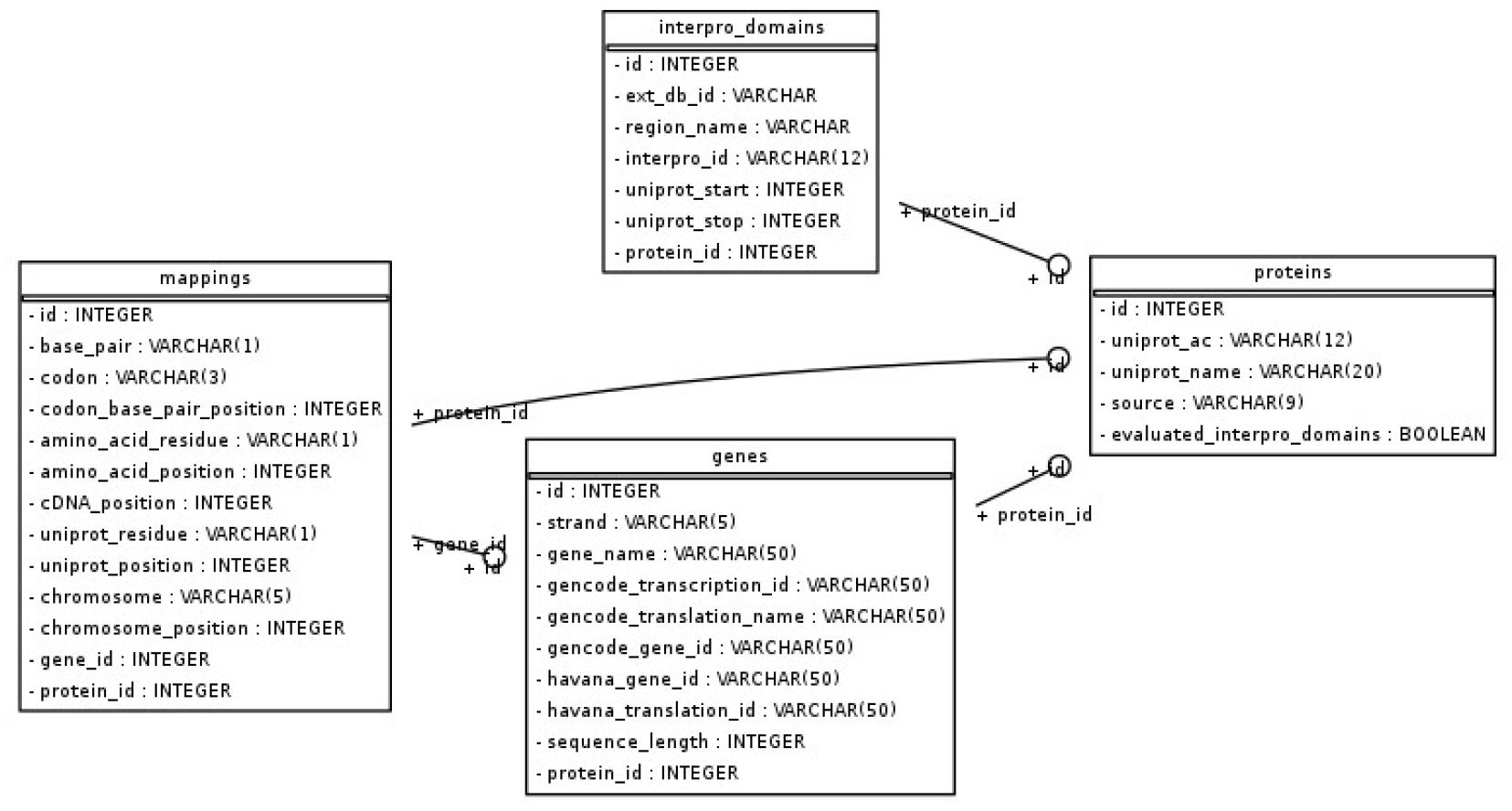
The internal database architecture for the MetaDome web server. The backbone of the MetaDome web server is a relational PostgreSQL database. This schematic representation of the tables present an overview of the relations in the data and what type of information is stored for each data entry. The table ‘mappings’ contain entries of a per-chromosome position (per gene) entries with codon to amino acid residue information. This is done for each of the nucleotides in a codon. The ‘genes’ table represent information from each of the transcripts present in the GENCODE database and the ‘proteins’ table correspond to global information on entries from the UniProtKB/Swiss-Prot databank. The ‘interpro_domains’ table can support any type of interpro domain, but for the purpose of MetaDome contain solely the Pfam protein domains. This database brings together genomic and protein positions that can be further used for annotation purposes.

